# Soil microbial communities are not altered by garlic mustard in recently invaded central Illinois forests

**DOI:** 10.1101/2020.12.19.423561

**Authors:** Joseph D. Edwards, Wendy H. Yang, Anthony C. Yannarell

**Author notes:** Corresponding author: Joseph Edwards.

## Abstract

The invasive forest plant garlic mustard (*Alliaria petiolata*) has been shown to alter soil microbial communities in the northeastern part of its invaded range in the United States, and this disruption of soil communities may contribute to its invasion success. However, garlic mustard allelochemistry can vary with invasion age, and it is not clear whether garlic mustard’s impacts on soil microbes are consistent over its invaded range. Here, we compare the composition and diversity of soil fungal, bacterial, and archaeal communities among garlic mustard present, absent, and removed treatments in replicated blocks across five forests in the midwestern United States with relatively young garlic mustard invasions (approximately 17-26 years old, with consistent management). We collected samples in May and August, corresponding to garlic mustard active and senescent life history stages. While soil fungal and bacterial/ archaeal communities (based on ITS2 region and 16S rRNA gene DNA sequencing, respectively) differed significantly between different blocks/ forests and over time, we found no significant effect of garlic mustard treatment on soil microbial community composition or the relative abundance of mycorrhizal, saprotrophic, or pathogenic fungal guilds. The lack of garlic mustard impacts on the soil microbial community in recently invaded central Illinois forests suggests that these well-documented impacts in the northeastern United States and in older invasions cannot necessarily be generalized across all environmental contexts.

## Introduction

Garlic mustard (*Alliaria petiolata*, Cavara and Grande) is a prominent invasive forest herb in North America (Rodgers et al. 2008), and it has been proposed that part of garlic mustard’s invasion success is due to novel interactions with soil microbiomes (Callaway et al. 2008). Garlic mustard produces secondary metabolites that reduce inoculum potential, activity, and biomass of mycorrhizal fungi (Anderson et al. 1996, Roberts and Anderson 2001). Because garlic mustard is a non-mycorrhizal plant, this assault on the mycorrhizal mutualists of its competitors has been posited to explain garlic mustard’s high invasion success (Stinson et al. 2006, Wolfe et al. 2008, Lankau et al. 2009). However, garlic mustard’s influence on microbial communities changes with invasion age because garlic mustard populations produce less allelochemicals over time (Lankau et al. 2009), thereby weakening its influence on soil microbes (Lankau et al. 2010). Furthermore, invasive plant effects on soil microbial communities (Arthur et al. 2012) and microbial processes (Portier et al. 2019) can also vary with plant phenology. Much of our understanding of garlic mustard impact of microbial communities is derived from studies on persistent invasions with high plant densities, mostly located in the north-eastern United States (Stinson et al. 2006, Wolfe et al. 2008, Anthony et al. 2017, Anthony et al. 2019, Burke et al. 2019a) and based on observations during garlic mustard’s active life period. Investigating the microbial impacts of younger, less dense garlic mustard populations in understudied regions of their invasion range and over different points in the plant’s life cycle can provide deeper insight into the microbial mechanisms by which garlic mustard promotes its invasion success.

Garlic mustard is primarily known for its allelopathic impacts on mycorrhizal fungi (Callaway et al. 2008), but garlic mustard invasions can also impact other members of the soil microbial community. For example, garlic mustard can promote fungal saprobes and pathogens (Anthony et al. 2017, Anthony et al. 2019), or decrease bacterial richness and abundance (Lankau et al. 2011). Promoting pathogens to which garlic mustard may be immune (Klironomos 2002) and altering decomposers to affect soil nutrient cycling (Hawkes et al. 2005, Hawkes et al. 2007) may potentially confer an advantage over native plants (Fraterrigo et al. 2011, Lee et al. 2018). However, garlic mustard effects on broader microbial communities are more poorly characterized than its effect on mycorrhizal communities (Cipollini and Cipollini 2016).

Invasive plant effects on soil microbial communities and their functions can vary with invasion age (Lankau 2011), plant phenology (Portier et al. 2019), and local soil properties (Li et al. 2006). After initial invasion, garlic mustard population densities increase rapidly then become relatively stable (Anderson et al. 1996, Nuzzo 1999). Because prior garlic mustard studies have focused primarily on older, more stable invasive populations, the effect of initial garlic mustard “booms” on soil microbiomes is not well understood. Many studies on microbial aspects of garlic mustard invasion also focus on early growing season, active life cycle plants, as these are most likely to have the greatest allelochemical production (Smith and Reynolds 2015). However, post-senescent or inactive invasive plants may still have significant impacts on microbial communities and processes (Arthur et al. 2012, Portier et al. 2019). Garlic mustard exhibits an early phenology relative to native competitors (Engelhardt and Anderson 2011), and so its effects on soil microbial communities may depend on its life stages or broader temporal changes associated with surrounding plants and ecosystem conditions (Saikkonen 2007, Bennett et al. 2013). Finally, local and landscape factors like disturbance regime, soil types, and surrounding plant communities also influence how quickly garlic mustard invasions spread and their population distribution patterns in invaded forests (Haines et al. 2018, Urbanowicz et al. 2019).

These factors suggest that garlic mustard’s influence on soil microbes could be variable across space (e.g. between plant populations in different forests) and time (e.g. between active vs. senescent garlic mustard plants).

Here, we asked the questions; 1) How does garlic mustard influence soil microbiomes in relatively young, low-density invasions; 2) Does this influence vary greatly within or between forests; and 3) How does this influence change over time between active and senescent garlic mustard plants? Using high-throughput DNA sequencing, we investigated how garlic mustard affects the community composition and species diversity of soil fungal and bacterial/archaeal communities between three garlic mustard treatments (present, absent, and removed) over two time points (May and August) across five hardwood forests in the midwestern United States. Our goal was to assess whether soils with garlic mustard present were more similar to each other than nearby reference soils (i.e., soils with garlic mustard absent or removed) across various spatial scales and as the growing season progressed. We used community composition to assess overall dissimilarity between samples, community diversity and evenness to assess whether garlic mustard was filtering surrounding microbiomes, and relative abundance of fungal functional guilds to assess the degree to which garlic mustard might be promoting its invasion success by suppressing mutualists, promoting pathogens, or altering nutrient cycling via saprotrophs.

## Methods

### Study Sites

We measured garlic mustard impacts on soil microbial communities across five forests in central Illinois: Brownfield Woods, Collins Pond Woods, Hart Woods, Richter Woods, and Trelease Woods. These sites are located within a 30 km north-south range and a 40 km east-west range. Mean annual temperature and mean annual precipitation ranged 10.0-11.1 °C and 889-977 mm, respectively, across all sites. The dominant soil series at each site were either silt loams or silty clay loams. Geographical coordinates, climate, soil type, and estimated invasion age for each site are reported in Table S1. Plant community composition is similar across all sites, with a “prairie grove” species mix with *Quercus, Fraxinus, Carya*, & *Acer* species dominating the canopies. The understories of these forests are made up of a diverse mix of tree seedlings, *Asimina triloba*, and other shrubs including *Sanicula, Laportea, Ageratina, Ribes*, and *Trillium* species. All sites are managed by the Illinois Natural History Survey (INHS; https://www.inhs.illinois.edu/research/natural-areas-uiuc). According to documentation from INHS managers, all of these sites began experiencing invasion between 1991 and 2000 (Table S1; S. Buck, personal communication). Site managers control garlic mustard populations through hand-pulling, herbicide application, and “weed whacking”, and as a result, garlic mustard densities at these sites are generally maintained below the high densities often associated with a decline in garlic mustard allelochemical production (Lankau et al. 2009, Lankau 2011). For all sites except for Collins Pond Woods, garlic mustard was restricted to small clusters of plants, with an average coverage of 30% of each m^2^ plot. Large portions of Collins Pond Woods were covered by garlic mustard plants, with an average coverage of 60% of each m^2^ plot.

### Study Design

In mid-March of 2017, we established four replicate 10-m^2^ experimental blocks in each forest. Each block contained 1-m^2^ plots for each of the following three treatments: garlic mustard present (present), garlic mustard absent (absent), and garlic mustard removed via hand-pulling (removed). We established the garlic mustard removed treatment as a control to account for unknown, meter-scale differences in soils within each block that might confound garlic mustard presence/absence in the other two plots, and the removal plots also allowed us to determine if live garlic mustard plants are required to see their effects on soil microbes. We collected soil samples in May and August, corresponding with the rosette and senescent life stages of garlic mustard, respectively. Soil samples were collected from 0-10 cm depth from the soil surface using a 2 cm diameter hand corer. There was very little, if any, development of organic horizons in any of our plots, so all samples consisted primarily of mineral soils. Four soil cores were collected from each 1 m^2^ treatment plot, composited, and homogenized by hand in the field. On the same day as collection, the soil samples were transported to the University of Illinois at Urbana-Champaign and frozen at −20°C for later analysis.

### DNA extraction, sequencing, identification, and assignments

We used high-throughput DNA sequencing to determine the impact of garlic mustard on microbial community structure as well as the relative abundance of key functional groups. DNA was extracted from 500 mg of freeze-dried soil using the FastDNA SPIN Kit for Soils (MP Biomedicals, Santa Ana, USA). The extracts were purified using cetyl trimethyl ammonium bromide (CTAB) as described in Edwards et al. (2018). DNA extracts were submitted to the Roy J. Carver Biotechnology Center at the University of Illinois at Urbana-Champaign for Fluidigm amplification (Fluidigm, San Francisco, USA) and Illumina sequencing (Illumina, San Diego, USA). We assessed fungal communities via the *ITS2* gene region using ITS3 and ITS4 (Ihrmark et al. 2012) primers to amplify DNA and sequence amplicons via MiSeq bulk 2 x 250 bp V2. As we were interested in the broad fungal community effect of garlic mustard invasion, we chose to use a universal ITS2 barcode region (Schoch et al. 2012). We assessed bacterial and archaeal communities via the bacterial and archaeal *16S* rRNA genes. The samples were amplified using V4_515 forward and V4_806 reverse primers and amplicons were sequences via MiSeq bulk 2 x 250 bp V2. Primer sequences and references can be found in Table S2. Sequence data is publicly available from the NCBI SRA database under accession number PRJNA690109.

We used the DADA2 pipeline (Callahan et al. 2016) for bioinformatic processing to produce amplicon sequence variants (ASVs) from the sequencing data. Briefly, quality filtering, denoising, mering forward and reverse reads, and removing chimeric sequences was performed using recommended parameters for *16S* and *ITS* (Callahan et al. 2020) genes. For *16S* processing one sample did not pass quality filtering (for a total of 119 samples), and for *ITS* processing two samples did not pass quality filtering (for a total of 118 samples). We did not cluster sequence variants (Glassman and Martiny 2018) prior to using the default DADA2 classifier (Wang et al. 2007) to assign taxonomy based on reference sequences from the UNITE database (Kõljalg et al. 2013). We relativized, rather than rarified (Gloor et al. 2017), our final count numbers via a Hellinger transformation (Legendre and Gallagher 2001) prior to further statistical analysis. Data used for analysis consisted of 9362 fungal, 14618 bacterial, and 152 archaeal ASVs. Due to the relatively low number of archaeal sequence variants, and because bacterial and archaeal sequences were derived from the same primer sets, we analyzed bacterial and archaeal communities together. We also compared the relative abundance and species richness of saprotrophic, pathogenic, and mycorrhizal functional guilds among garlic mustard treatments and between sampling dates using FUNGuild (Nguyen et al. 2016) to assign functional guild to taxa. Assignments with “probable” or “highly probable” confidence scores were included and in cases where taxa were assigned to more than one guild, mycorrhizal classification was given higher priority than pathogenic, which was given higher priority than saprotrophic (Smith et al. 2021).

### Statistical Analysis

All statistical analyses were performed in the R statistical environment (Team 2013). Overall patterns of beta diversity and community structure were analyzed with the VEGAN package (Oksanen et al. 2010). To determine differences in community composition based on treatment, sampling date, and the interaction between treatment and sampling date, we performed a permutational analysis of variance (PERMANOVA) with the *adonis* function on Hellinger-transformed relativized ASV count numbers. We added location (forest or block) as a term in the *adonis* model, rather than stratifying on these terms, in order to estimate the spatial contribution to the total variance in microbial communities; comparison of these models with those that stratified on location instead did not produce different results with respect to treatment, sampling date, or their interaction (Table S3). We also assessed differences in community composition and heterogeneity visually using nonmetric multidimensional scaling (NMDS) ordinations on Manhattan dissimilarity of relativized ASV count numbers; this distance measure was chosen using the *rankindex()* function in *vegan*. We assessed alpha diversity as species diversity (Shannon’s *H*) and evenness (*H*/log(species #)) using the *diversity()* function in *vegan*. Differences among garlic mustard treatments, sampling dates, and their interaction in the species richness and evenness, as well as relative abundance and richness of functional guilds, were determined using a linear-mixed model with the LMER package (De Boeck et al. 2011), mycorrhizal abundance and richness values were log-transformed to meet assumptions of normality. In this model, we used treatment, sampling date, and their interaction as fixed effects and block nested within forest as a random effect. Statistical significance was assessed as p < 0.05.

## Results

### Soil microbial community composition

Sampling date and forest significantly influenced fungal and bacterial/archaeal community composition, though neither garlic mustard treatment nor its interaction with sampling date had a significant effect (Table 1). Across all samples, forest was responsible for 11.5% of variation in fungal communities, while within forests the variation attributable to block increased to between 21%-27% (p<0.001, Table 1, Figure 1C). For bacterial and archaeal communities forest was responsible for 15.7% of variation among all samples and between 28%-46% for block within each forest (p<0.001, Table 1, Figure 1F). Sampling date accounted for 2.2% of variation in fungal communities among all forests (p<0.001, Table 1, Figure 1B) and between 4.9% and 12.9% (p<0.05, Table 1, Figure 2F-J) in all forests except Collins Pond where it did not have a significant effect. For bacterial and archaeal communities, sampling date was responsible for 4.2% of overall variation among all samples (p<0.001, Table 1, Figure 1E) and between 8.3%-10.2% (p<0.006, Table 1, Figure 3F-J) in all forests except Hart Woods where it did not have a significant effect. Garlic mustard treatment did not significantly affect overall fungal communities across all forests or within any forest (Figure 1A, Figure 2A-E). Similarly, garlic mustard treatment also had no significant effect on bacterial and archaeal communities across all forests or within any forest (Figure 1D, Figure 3A-E).

**Figure 1.**
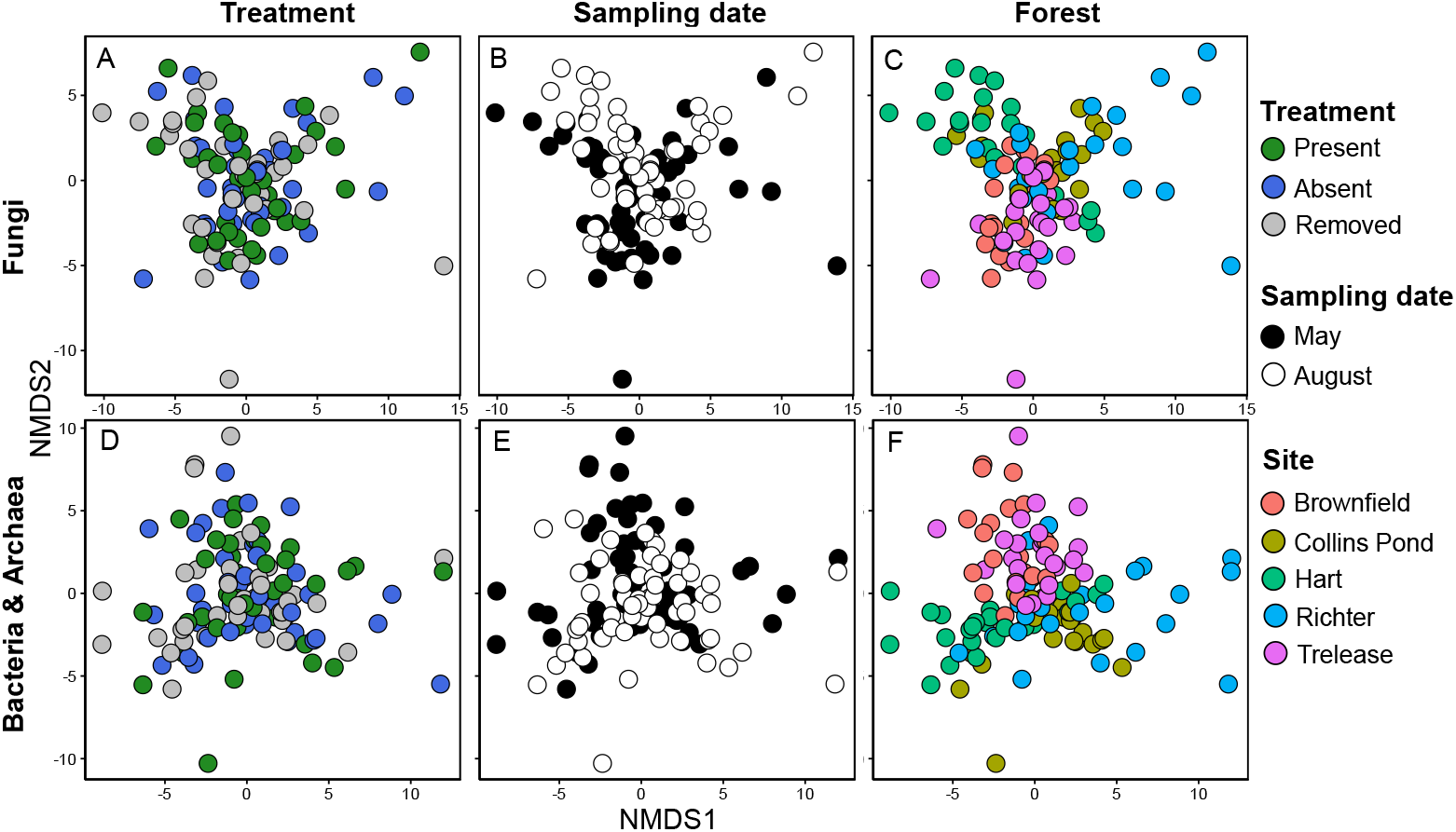
Fungal (ABC, Stress = 0.15) and Bacterial/Archaeal (DEF, Stress = 0.07) community composition expressed as nonmetric multidimensional scaling axes depicting Manhattan dissimilarities of relativized count numbers for ASVs within each sample (circle). Panels A & D depict differences between garlic mustard invasion status (treatment), panels B & E depict differences between sampling dates, and panels C and F depict differences between forests. Each panel includes data across all garlic mustard treatments, sampling dates, and forests.

**Figure 2.**
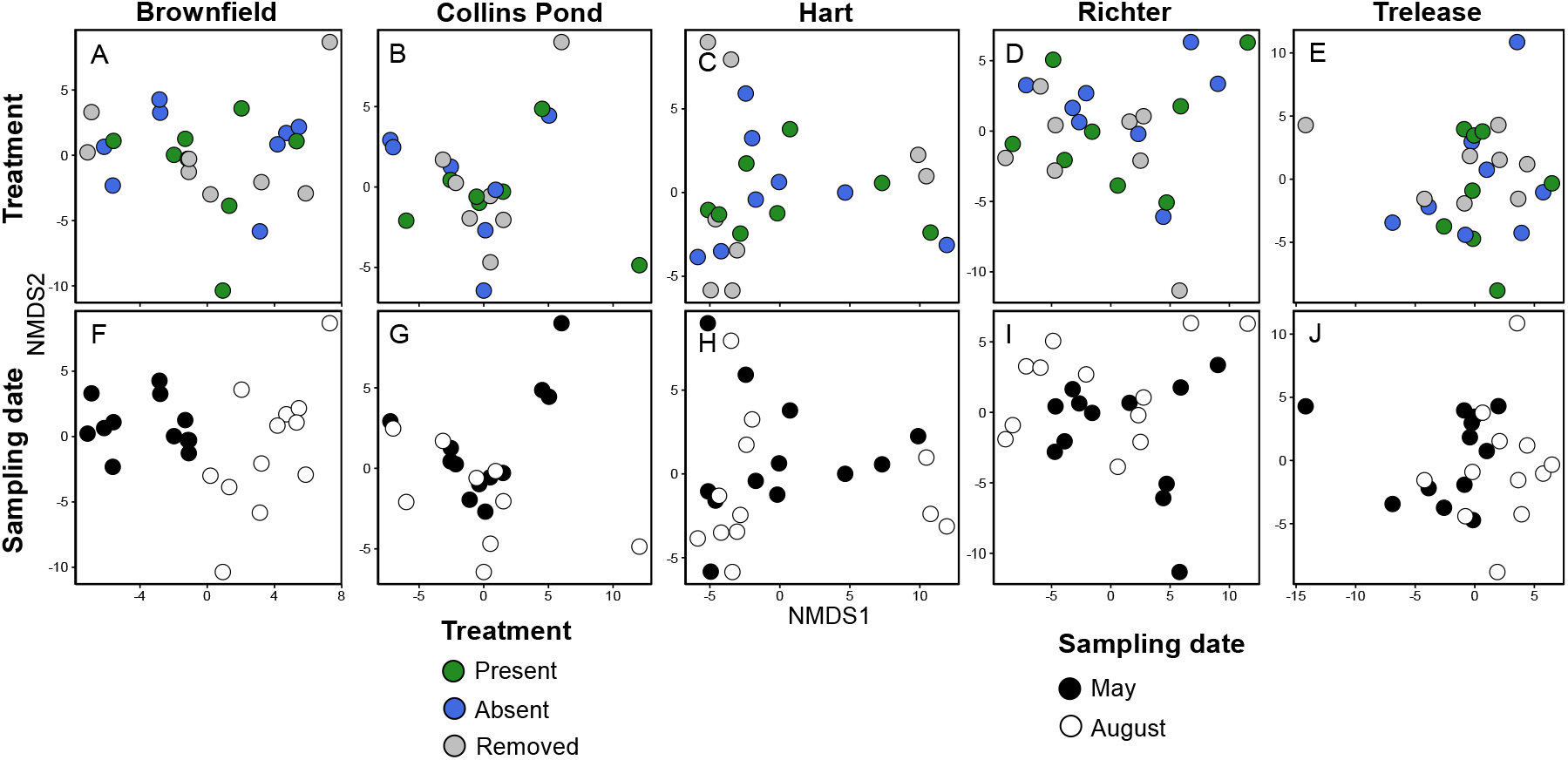
Individual forest ordinations of fungal community composition expressed as nonmetric multidimensional scaling axes depicting Manhattan dissimilarities of relativized count numbers for ASVs within each sample (circle). Panels A-E depict differences between garlic mustard invasion status while panels F-J depict differences between sampling dates, with each panel only containing data from the forest listed above each column. Ordination stress ranged from 0.06-0.10.

**Figure 3.**
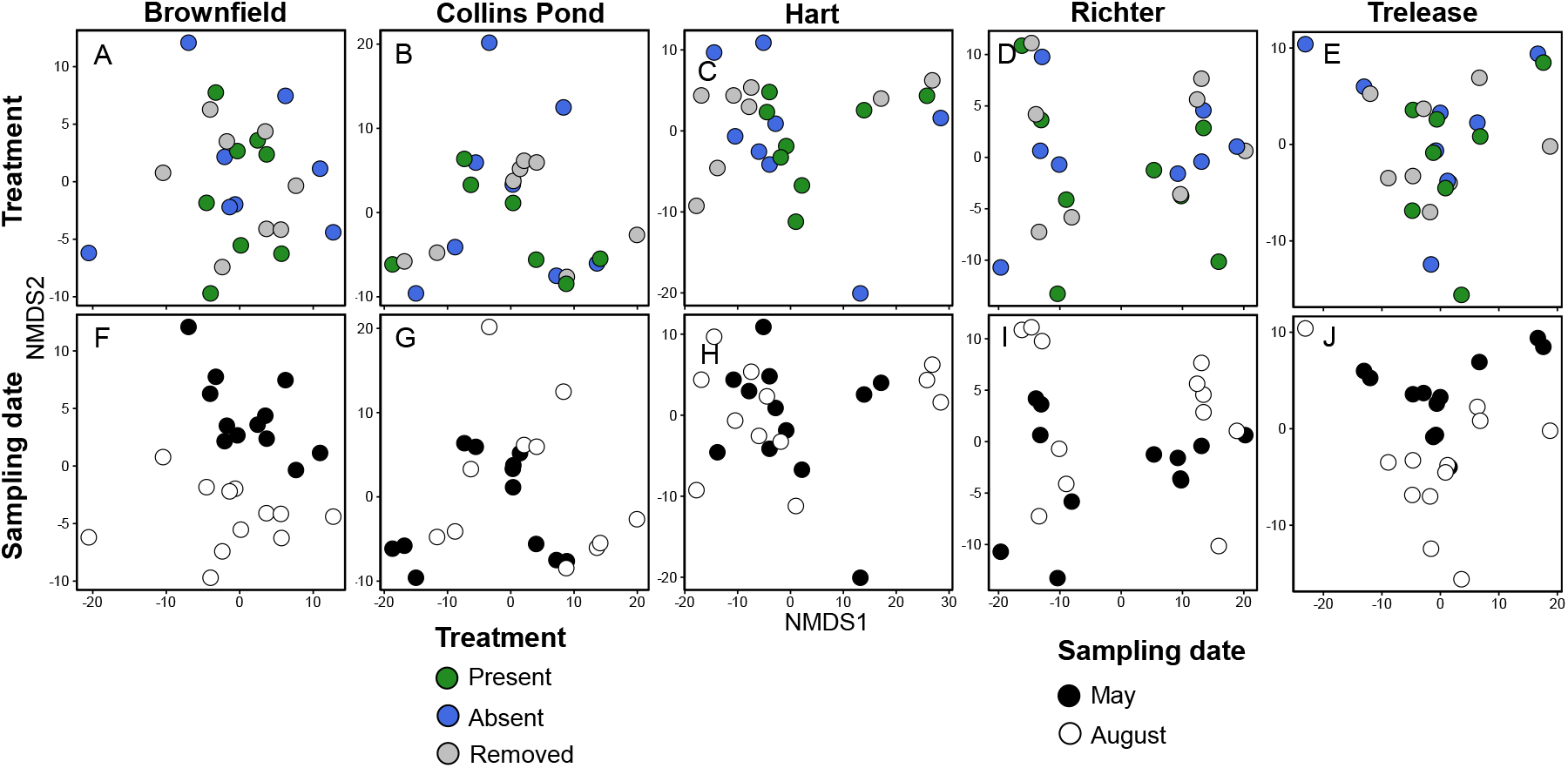
Individual forest ordinations of bacterial and archaeal community composition expressed as nonmetric multidimensional scaling axes depicting Manhattan dissimilarities of relativized count numbers for ASVs within each sample (circle). Panels A-E depict differences between garlic mustard invasion status while panels F-J depict differences between sampling dates, with each panel only containing data from the forest listed above each column. Ordination stress ranged from 0.03-0.08.

**Table 1.** PERMANOVA results for fungal and bacterial/archaeal community composition across all forests. Bolded p-values highlight significant effects at p < 0.05.

| Forest | Community | Main Fixed Effect | F-value | R<sup>2</sup> | p-value |
| --- | --- | --- | --- | --- | --- |
| All | Fungi | Treatment | 0.884 | 0.013 | 0.932 |
|  |  | Sampling date | 2.665 | 0.022 | **0.001** |
|  |  | Treatment*Sampling date | 0.74 | 0.012 | 1.000 |
|  |  | Forest | 3.678 | 0.115 | **0.001** |
|  | Bacteria & Archaea | Treatment | 0.900 | 0.013 | 0.652 |
|  |  | Sampling date | 5.843 | 0.042 | **0.001** |
|  |  | Treatment*Sampling date | 0.648 | 0.009 | 1.000 |
|  |  | Forest | 5.5004 | 0.157 | **0.001** |
| Brownfield | Fungi | Treatment | 0.925 | 0.650 | 0.705 |
|  |  | Sampling date | 3.670 | 0.129 | **0.001** |
|  |  | Treatment*Sampling date | 0.893 | 0.062 | 0.748 |
|  |  | Block | 2.043 | 0.216 | **0.001** |
|  | Bacteria & Archaea | Treatment | 1.039 | 0.068 | 0.342 |
|  |  | Sampling date | 3.118 | 0.102 | **0.001** |
|  |  | Treatment*Sampling date | 0.835 | 0.055 | 0.873 |
|  |  | Block | 2.863 | 0.282 | **0.001** |
| Collins Pond | Fungi | Treatment | 1.022 | 0.090 | 0.411 |
|  |  | Sampling date | 1.183 | 0.051 | 0.137 |
|  |  | Treatment*Sampling date | 0.865 | 0.075 | 0.907 |
|  |  | Block | 2.040 | 0.265 | **0.001** |
|  | Bacteria & Archaea | Treatment | 1.038 | 0.062 | 0.363 |
|  |  | Sampling date | 3.344 | 0.100 | **0.001** |
|  |  | Treatment*Sampling date | 0.929 | 0.056 | 0.577 |
|  |  | Block | 4.014 | 0.361 | **0.001** |
| Hart | Fungi | Treatment | 1.061 | 0.076 | 0.265 |
|  |  | Sampling date | 1.534 | 0.055 | **0.013** |
|  |  | Treatment*Sampling date | 0.880 | 0.063 | 0.814 |
|  |  | Block | 2.548 | 0.272 | **0.001** |
|  | Bacteria & Archaea | Treatment | 1.337 | 0.082 | 0.108 |
|  |  | Sampling date | 1.725 | 0.053 | 0.066 |
|  |  | Treatment*Sampling date | 0.786 | 0.049 | 0.839 |
|  |  | Block | 3.779 | 0.351 | **0.001** |
| Richter | Fungi | Treatment | 1.072 | 0.077 | 0.231 |
|  |  | Sampling date | 1.366 | 0.049 | **0.047** |
|  |  | Treatment*Sampling date | 0.903 | 0.0645 | 0.807 |
|  |  | Block | 2.5077 | 0.270 | **0.001** |
|  | Bacteria & Archaea | Treatment | 1.064 | 0.051 | 0.381 |
|  |  | Sampling date | 3.457 | 0.083 | **0.006** |
|  |  | Treatment*Sampling date | 0.870 | 0.042 | 0.570 |
|  |  | Block | 6.452 | 0.464 | **0.001** |
| Trelease | Fungi | Treatment | 1.001 | 0.074 | 0.453 |
|  |  | Sampling date | 1.941 | 0.072 | **0.001** |
|  |  | Treatment*Sampling date | 0.831 | 0.061 | 0.961 |
|  |  | Block | 2.140 | 0.238 | **0.001** |
|  | Bacteria & Archaea | Treatment | 1.132 | 0.072 | 0.225 |
|  |  | Sampling date | 3.196 | 0.101 | **0.001** |
|  |  | Treatment*Sampling date | 0.843 | 0.054 | 0.827 |
|  |  | Block | 3.119 | 0.297 | **0.001** |

**Table 2.** ANOVA results for main effects on the species diversity (Shannon’s *H*) and evenness (*H*/log(species #)). Block nested within site was included as a random effect in all models. Bolded p-values highlight significant effects at p < 0.05.

| Community | Main Fixed Effect | Diversity |  | Evenness |  |
| --- | --- | --- | --- | --- | --- |
|  |  | F-value | p-value | F-value | p-value |
| **Fungi** | Treatment | 0.274 | 0.761 | 0.059 | 0.943 |
|  | Sampling date | 0.128 | 0.721 | 34.82 | **<0.001** |
|  | Treatment*Sampling date | 3.188 | **0.045** | 1.178 | 0.312 |
| **Bacteria & Archaea** | Treatment | 2.417 | 0.095 | 2.404 | 0.096 |
|  | Sampling date | 2.599 | 0.110 | 41.014 | **<0.001** |
|  | Treatment*Sampling date | 0.597 | 0.552 | 6.113 | **0.003** |

### Diversity and evenness

Garlic mustard presence did not significantly affect microbial diversity or evenness relative to control or removed plots, though fungal diversity increased from May to August corresponding with plant senescence. The interaction between garlic mustard treatment and time had a significant effect on fungal diversity (*F_2,112_*=3.18, p=0.045), with diversity increasing in garlic mustard present plots from May to August (Figure 4A). Fungal evenness increased significantly from May to August (*F_1,112_*=34.82, p<0.001), though garlic mustard treatment had no significant effect (Figure 4B). There were no significant effects or interactions of garlic mustard treatment and time on bacterial and archaeal diversity (Figure 4C), though garlic mustard treatment was marginally significant (p=0.095). Bacterial and archaeal evenness significantly increased from May to August (*F_1,113_*=41.014, p<0.001), and this effect was most pronounced in garlic mustard present plots (interaction *F_2,113_*=6.113, p=0.003, Figure 4C).

**Figure 4.**
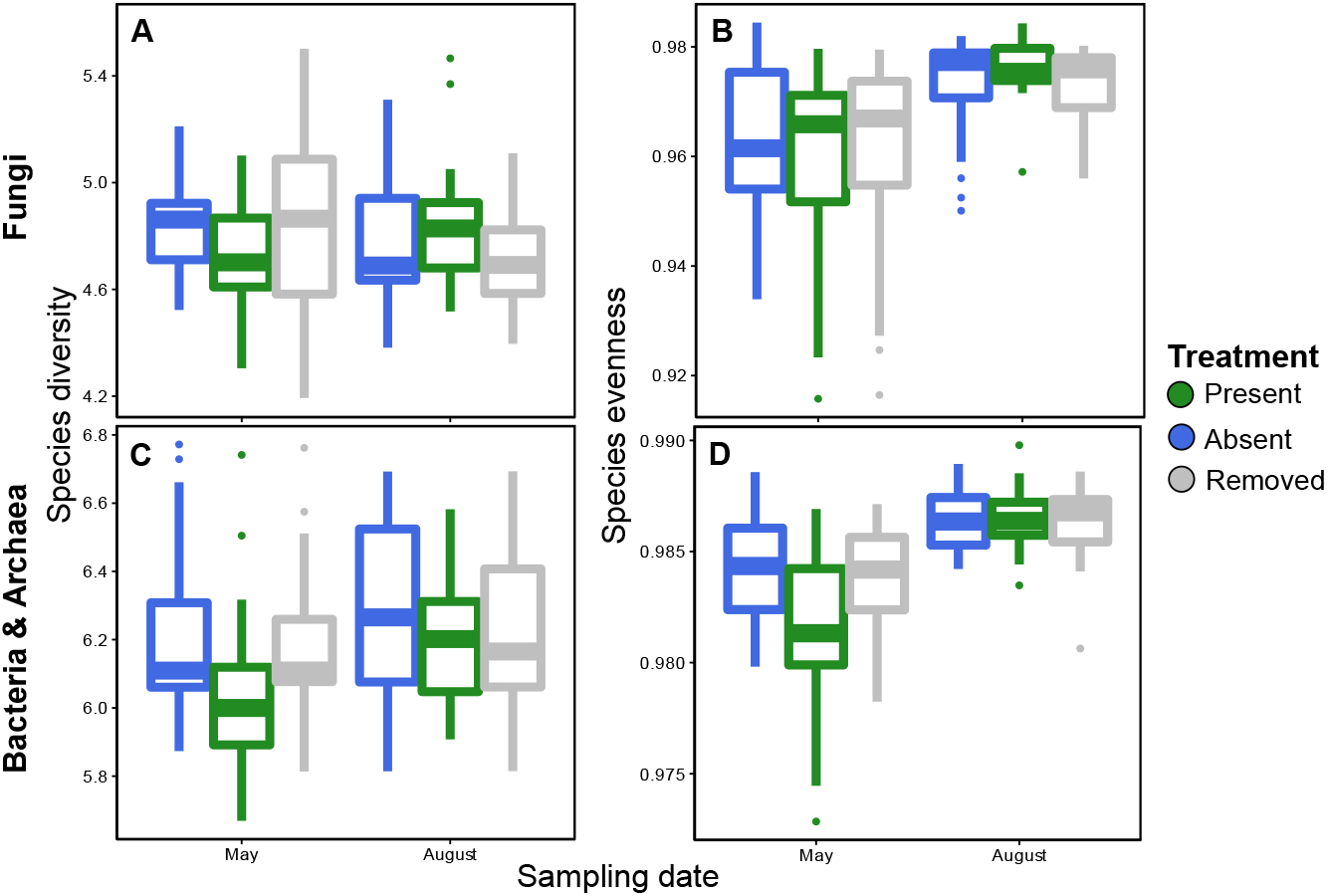
Diversity indices for fungal (AB) and bacterial/archaeal (CD) communities. Panels A and C show species diversity, expressed as Shannon’s *H*, and panels B and D show species evenness, expressed as *H*/log(species #).

### Functional guilds

Garlic mustard treatment had no significant effect on the relative abundance or species richness of fungal saprotrophic, pathogenic, and mycorrhizal functional guilds (Table S4, Figure S1). Mycorrhizal relative abundance and richness increased significantly from May to August (relative abundance: *F_1,112_*=29.762, p < 0.001; richness: *F_1,112_*=26.202, p < 0.001, Figure S1CF). Fungal pathogen richness decreased significantly from May to August (*F_1,112_*=7.425, p=0.007), though this effect was most pronounced in the garlic mustard removed plots (interaction: *F_2,112_*=4.928, p=0.009, Figure S1E).

## Discussion

We found extensive spatial variation of microbial community composition both within and between forests, but we did not detect consistent or strong microbial patterns related to garlic mustard presence at these sites. Despite long known (Vaughn and Berhow 1999, Roberts and Anderson 2001) and widely reported effect of garlic mustard on soil microbiomes (Stinson et al. 2006, Callaway et al. 2008, Rodgers et al. 2008, Wolfe et al. 2008, Pringle et al. 2009, Anderson et al. 2010, Anthony et al. 2017), garlic mustard appears to have minimal impacts on the soil microbiome of these recently invaded forests of central Illinois. Further, we also did not find support for the “mycorrhizal suppression” that is often reported with garlic mustard invasions (Stinson et al. 2006, Wolfe et al. 2008), nor any direct impact on other fungal functional guilds that have been more recently reported (Anthony et al. 2017, Duchesneau et al. 2021). While soil microbial communities at our sites did change significantly between our May and August sampling dates, we were unable to detect significant garlic mustard effects on soil microbial communities during either the active or senescent garlic mustard phenological stages. Overall, our study suggests that caution should be taken in generalizing microbially-mediated invasion mechanisms for garlic mustard across all potential ecoregions, ecosystems, and invasion stages.

Garlic mustard’s minimal impact on soil microbial communities in our study could potentially be due to regional or local soil characteristics of our sites. Many of the soils in our study had higher clay contents than the characteristically sandy soil of more northern and eastern North American forests invaded by garlic mustard (Table S1). Allelochemicals produced by garlic mustard plants could have bound to these clay particles (Blair et al. 2005), potentially ameliorating their inhibitory effects on mycorrhizal fungi (Cantor et al. 2011). While some of the earliest evidence for garlic mustard’s mycorrhizal suppression was also found in central Illinois soils (Anderson et al. 1996, Roberts and Anderson 2001), these studies found garlic mustard decreased mycorrhizal inoculum potential, which could occur without a concomitant change in microbial community composition. The possibility of certain soils mediating garlic mustard’s allelopathy could have ramifications for future garlic mustard invasion pattens, though further investigation of the relationship between allelochemical concentrations and microbial responses across different soil types will be required to support this pattern.

The age of garlic mustard populations in our study may also have influenced their impacts on soil microbial communities. Microbial community suppression by garlic mustard peaks when invasive populations are around 40 years old (Lankau 2011), which may represent a potential ecological filter on microbial communities in garlic mustard invaded soils. According to management records from these sites all garlic mustard population were <26 years old and, due to land management, with low garlic mustard plant density characteristic of much younger invasions. We found weak evidence of decreased soil microbial diversity in garlic mustard present plots, particularly in May when the plants were active, which could possibly be an indicator of ongoing microbiome manipulation by garlic mustard populations. As the garlic mustard populations we observed in this study have not yet reached the age associated with peak allelochemical production (Lankau 2011), the microbial filtering effect of this allelochemical peak may not yet be fully present. Ecological filtering associated with invasion age or density could explain differences in our results, particularly mycorrhizal suppression, from studies conducted on older, higher density garlic mustard populations (Anthony et al. 2019, Burke et al. 2019b). Direct comparisons between study settings are needed to elucidate whether these differences are due to invasion age or regional differences in ecosystem characteristics.

The influence of spatial variation on microbial communities is often considered a key factor driving community composition (Fierer and Lennon 2011). These patterns, as we saw in this study, are often similar (or even stronger) at meter to decameter scales than at regional to global scales (Ramirez et al. 2014). The many factors that promote this variation (e.g. temperature, moisture, pH, soil chemistry, trophic interactions, plant and animal communities, etc.) necessitates the need for contextualizing extrinsic impacts on microbial communities, like those of an invasive plant, with their neighboring microbiomes. Experimental designs that account for this variation by comparing the impacts of invasive species with closely situated control soils, like the hierarchical block-with-forest design we used here, should be utilized in future investigations to ensure patterns are consistent and generalizable across a large area. Temporal variation also played a significant role in structuring microbial communities, particularly the increase in mycorrhizal abundance and richness over the growing season. The impacts of time on microbial community composition is less well understood than that of space (Carini et al. 2020). Temporal factors are likely to be more influenced by short-term patterns, like weather or plant phenology, than spatial factors potentially making them more susceptible to future global change (Balser et al. 2010). Further research on short-term temporal patterns, like phenology or warming, in invasive species-microbiome interactions (Anthony et al. 2020) is needed to further understand how microbial communities respond to invasive species and their environment over time. Improving understanding of spatial and temporal variation in microbially mediated invasion dynamics could help predict the spread of future invasions and their responses to global change.

Overall, we present evidence that garlic mustard invasions may vary in their ability to manipulate microbiomes to promote their invasion success across ecoregions. This variation may be due to soil condition, invasion age, or some other factor. Invasive garlic mustard populations were still present in these forests despite their apparently limited microbial impacts. This finding suggests that garlic mustard populations are not solely reliant on microbial mechanisms (like “mycorrhizal suppression” to the degree we could measure it) to invade forest ecosystem. The lack of garlic mustard impacts on the soil microbial community in recently invaded central

Illinois forests suggests that these well-documented impacts in the northeastern United States and in older invasions cannot necessarily be generalized across all environmental contexts.

## Acknowledgments

We would like to thank the Illinois Natural History Survey and Grand Prairie Friends, particularly Jamie Ellis and Jeff Peyton, for allowing land access conducting this research and their gracious help in finding garlic mustard populations. We greatly appreciate the assistance from Ally Cook, Alex Kent, Alex Krichels, Elle Lucadamo, Nikki Snyder, Xiangyu Zhang, Joseph Schlosser, Alonso Favela, and Gabe Price for their help in establishing treatments and sample collection. This work was supported by the National Institute of Food and Agriculture, U.S. Department of Agriculture, McIntire Stennis program under accession number 1011139 to ACY. JDE was also supported by the Cooperative State Research, Education, and Extension Service, US Department of Agriculture, under project number ILLU 875-952. We are also grateful to Adam Davis and James Dalling for comments on earlier versions of this manuscript.

## Supplementary Material

**Table S1.** Study site locations, climate, soil type, and year garlic mustard was first reported. Climate and soil data was obtained from the USDA Web Soil Survey (https://websoilsurvey.sc.egov.usda.gov/) and accessed on [12/06/2020], climate data was averaged between years 2002-2020. Garlic mustard report year was provided by personal communication from S. Buck, the natural areas manager at these sites.

| Site name | Latitude/<br>Longitude<br>(°N, °W) | Mean annual<br>temperature<br>(°C) | Mean annual<br>precipitation<br>(mm) | Dominant soil series | Year garlic<br>mustard first<br>reported |
| --- | --- | --- | --- | --- | --- |
| **Brownfield**<br>**Woods** | 40.14, -88.16 | 10.0 | 965 | Drummer silty clay<br>loam | 1995 |
| **Collins Pond**<br>**Woods** | 40.13, -88.03 | 10.5 | 990 | Raub silt loam and<br>Xenia silt loam | 1997 |
| **Hart Woods** | 40.22, -88.35 | 10.0 | 889 | Ambraw silty clay<br>loam and Martinsville<br>silt loam | 1995 |
| **Richter**<br>**Woods** | 40.09, -87.81 | 10.0 | 914 | Birkbeck silt loam and<br>Strawn silt loam | 2000 |
| **Trelease**<br>**Woods** | 40.13, -88.14 | 10.0 | 965 | Flanagan silt loam and<br>Drummer silty clay<br>loam | 1991 |

**Table S2.**
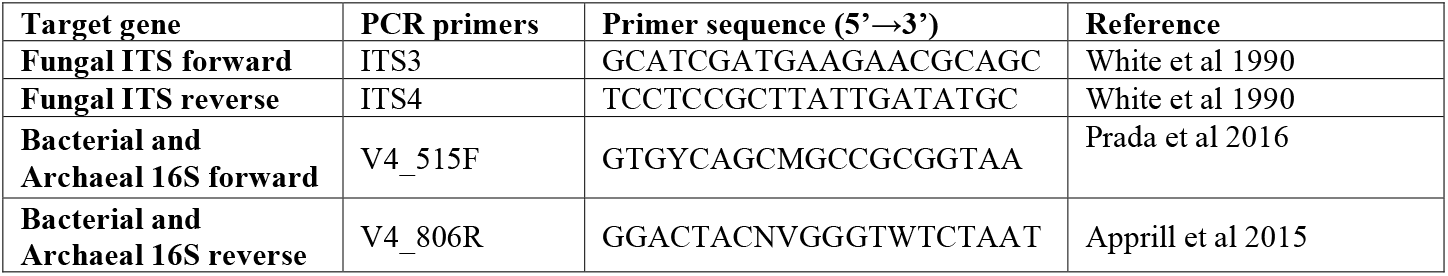
Target genes, PCR primer names, primer sequences, and references for genes sequences in fungal and bacterial/ archaeal community analysis.

**Table S3.** PERMANOVA test on fungal and bacterial/ archaeal community composition across all forests with forest stratified in the model. Bolded p-values highlight significant effects at p < 0.05.

| Forest | Community | Main Fixed Effect | F-value | R<sup>2</sup> | p-value |
| --- | --- | --- | --- | --- | --- |
| All | Fungi | Treatment | 0.806 | 0.013 | 0.974 |
|  |  | Sampling date | 2.613 | 0.022 | **0.001** |
|  |  | Treatment*Sampling date | 0.683 | 0.011 | 1.000 |
|  | Bacteria & Archaea | Treatment | 0.776 | 0.013 | 0.717 |
|  |  | Sampling date | 5.040 | 0.417 | **0.001** |
|  |  | Treatment*Sampling date | 0.555 | 0.009 | 1.000 |

**Table S4.** Table 2. ANOVA results for main effects on the relative abundance (% read #) and species richness (#) of saprotrophic, pathogenic, and mycorrhizal fungi. Block nested within site was included as a random effect in all models. Bolded p-values highlight significant effects at p < 0.05.

| Measure | Main Fixed Effect | Saprotroph |  | Pathogen |  | Mycorrhizal |  |
| --- | --- | --- | --- | --- | --- | --- | --- |
|  |  | F-value | p-value | F-value | p-value | F-value | p-value |
| Relative abundance | Treatment | 0.685 | 0.506 | 0.055 | 0.947 | 0.700 | 0.499 |
|  | Sampling date | 3.032 | 0.085 | 0.449 | 0.504 | 29.762 | **<0.001** |
|  | Treatment*Sampling date | 1.259 | 0.289 | 1.169 | 0.315 | 0.048 | 0.953 |
| Richness | Treatment | 1.186 | 0.310 | 0.120 | 0.887 | 1.318 | 0.273 |
|  | Sampling date | 2.880 | 0.093 | 7.425 | **0.007** | 26.202 | **<0.001** |
|  | Treatment*Sampling date | 1.230 | 0.297 | 4.928 | **0.009** | 0.190 | 0.827 |

**Figure S1.**
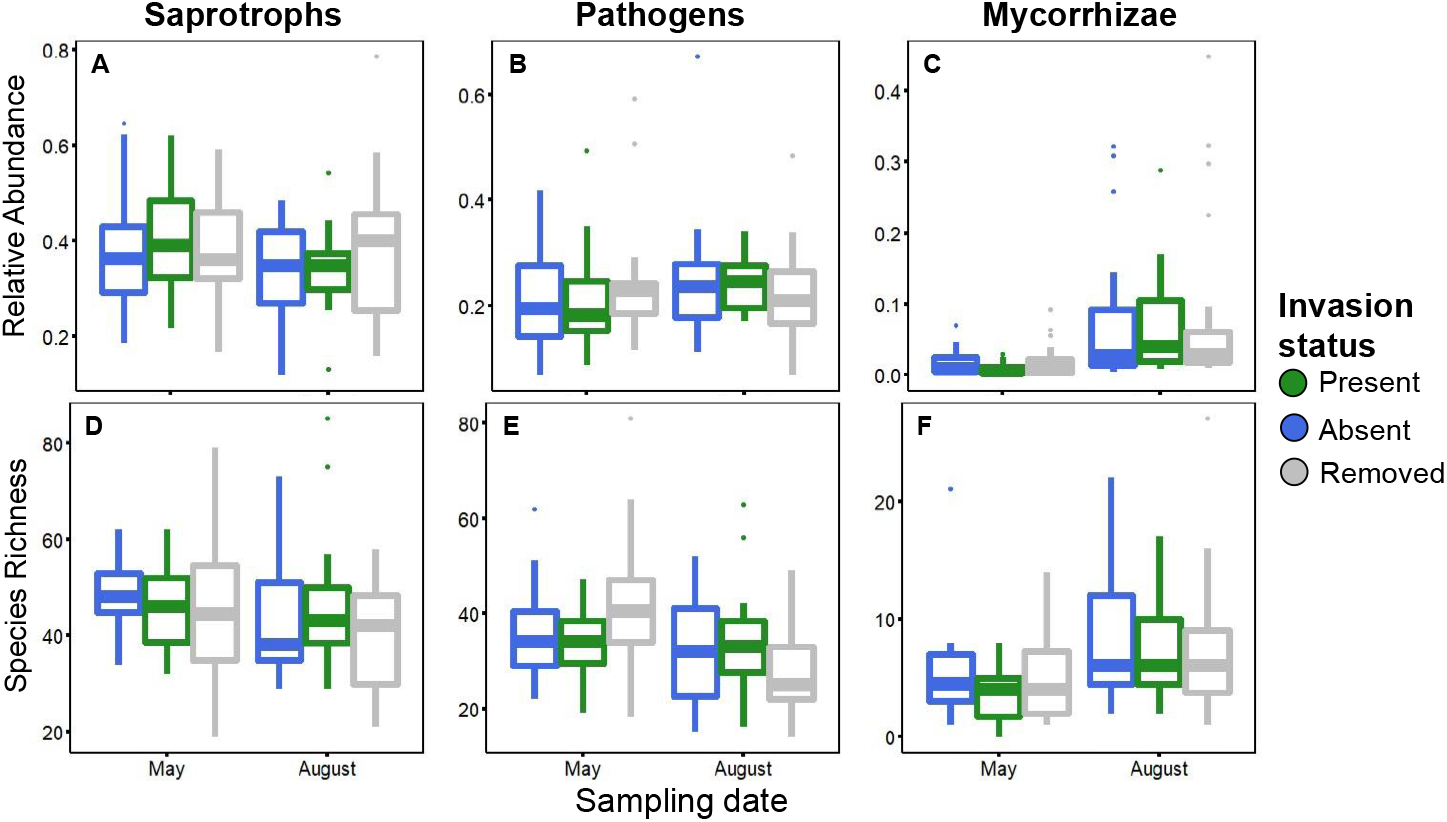
Relative abundance (A-C) and species richness (D-E) of saprotrophic (AD), pathogenic (BE), and mycorrhizal (CF) functional guilds

## References

Anderson, R. C., M. R. Anderson, J. T. Bauer, M. Slater, J. Herold, P. Baumhardt, and V. Borowicz. 2010. Effect of removal of garlic mustard (*Alliaria petiolata*, Brassicaeae) on arbuscular mycorrhizal fungi inoculum potential in forest soils. The Open Ecology Journal 3:41–47.

Anderson, R. C., S. S. Dhillion, and T. M. Kelley. 1996. Aspects of the ecology of an invasive plant, garlic mustard (*Alliaria petiolata*), in central Illinois. Restoration Ecology 4:181–191.

Anthony, M., S. Frey, and K. Stinson. 2017. Fungal community homogenization, shift in dominant trophic guild, and appearance of novel taxa with biotic invasion. Ecosphere 8:e01951.

Anthony, M., K. Stinson, A. Trautwig, E. Coates-Connor, and S. Frey. 2019. Fungal communities do not recover after removing invasive *Alliaria petiolata* (garlic mustard). Biological invasions 21:3085–3099.

Anthony, M. A., K. A. Stinson, J. A. Moore, and S. D. Frey. 2020. Plant invasion impacts on fungal community structure and function depend on soil warming and nitrogen enrichment. Oecologia:1–14.

Apprill, A., S. McNally, R. Parsons, and L. Weber. 2015. Minor revision to V4 region SSU rRNA 806R gene primer greatly increases detection of SAR11 bacterioplankton. Aquatic Microbial Ecology 75:129–137.

Arthur, M. A., S. R. Bray, C. R. Kuchle, and R. W. McEwan. 2012. The influence of the invasive shrub, *Lonicera maackii*, on leaf decomposition and microbial community dynamics. Plant Ecology 213:1571–1582.

Balser, T. C., J. L. Gutknecht, and C. Liang. 2010. How will climate change impact soil microbial communities? Pages 373–397 Soil microbiology and sustainable crop production. Springer.

Bennett, A. E., T. J. Daniell, M. Öpik, J. Davison, M. Moora, M. Zobel, M.-A. Selosse, and D. Evans. 2013. Arbuscular mycorrhizal fungal networks vary throughout the growing season and between successional stages. PLoS One 8:e83241.

Blair, A. C., B. D. Hanson, G. R. Brunk, R. A. Marrs, P. Westra, S. J. Nissen, and R. A. Hufbauer. 2005. New techniques and findings in the study of a candidate allelochemical implicated in invasion success. Ecology Letters 8:1039–1047.

Burke, D. J., S. R. Carrino-Kyker, A. Hoke, S. Cassidy, L. Bialic-Murphy, and S. Kalisz. 2019a. Deer and invasive plant removal alters mycorrhizal fungal communities and soil chemistry: Evidence from a long-term field experiment. Soil Biol Biochemistry 128:13–21.

Burke, D. J., S. R. Carrino-Kyker, A. Hoke, S. Cassidy, L. Bialic-Murphy, and S. Kalisz. 2019b. Deer and invasive plant removal alters mycorrhizal fungal communities and soil chemistry: Evidence from a long-term field experiment. Soil Biology and Biochemistry 128:13–21.

Callahan, B., P. McMurdie, M. Rosen, A. Han, A. Johnson, and S. Holmes. 2020. DADA2 ITS pipeline workflow (1.8). Retrieved September 18:2020.

Callahan, B. J., P. J. McMurdie, M. J. Rosen, A. W. Han, A. J. A. Johnson, and S. P. Holmes. 2016. DADA2: high-resolution sample inference from Illumina amplicon data. Nature methods 13:581–583.

Callaway, R. M., D. Cipollini, K. Barto, G. C. Thelen, S. G. Hallett, D. Prati, K. Stinson, and J. Klironomos. 2008. Novel weapons: Invasive plant suppresses fungal mutualists in America but not in its native Europe. Ecology 89:1043–1055.

Cantor, A., A. Hale, J. Aaron, M. B. Traw, and S. Kalisz. 2011. Low allelochemical concentrations detected in garlic mustard-invaded forest soils inhibit fungal growth and AMF spore germination. Biological invasions 13:3015–3025.

Carini, P., M. Delgado-Baquerizo, E.-L. S. Hinckley, H. Holland-Moritz, T. E. Brewer, G. Rue, C. Vanderburgh, D. McKnight, and N. Fierer. 2020. Effects of spatial variability and relic DNA removal on the detection of temporal dynamics in soil microbial communities. Mbio 11:e02776–02719.

Cipollini, D., and K. Cipollini. 2016. A review of garlic mustard *(Alliaria petiolata*, Brassicaceae) as an allelopathic plant. J Torr Bot Soc 143:339–348.

De Boeck, P., M. Bakker, R. Zwitser, M. Nivard, A. Hofman, F. Tuerlinckx, and I. Partchev. 2011. The estimation of item response models with the lmer function from the lme4 package in R. J Stat Software 39:1–28.

Duchesneau, K., A. Golemiec, R. I. Colautti, and P. M. Antunes. 2021. Functional shifts of soil microbial communities associated with Alliaria petiolata invasion. Pedobiologia 84:150700.

Edwards, J. D., C. M. Pittelkow, A. D. Kent, and W. H. Yang. 2018. Dynamic biochar effects on soil nitrous oxide emissions and underlying microbial processes during the maize growing season. Soil Biology and Biochemistry 122:81–90.

Engelhardt, M. J., and R. C. Anderson. 2011. Phenological niche separation from native species increases reproductive success of an invasive species: *Alliaria petiolata* (Brassicaceae)– garlic mustard. J Torr Bot Soc 138:418–433.

Fierer, N., and J. T. Lennon. 2011. The generation and maintenance of diversity in microbial communities. American Journal of Botany 98:439–448.

Fraterrigo, J. M., M. S. Strickland, A. D. Keiser, and M. A. Bradford. 2011. Nitrogen uptake and preference in a forest understory following invasion by an exotic grass. Oecologia 167:781.

Glassman, S. I., and J. B. Martiny. 2018. Broadscale ecological patterns are robust to use of exact sequence variants versus operational taxonomic units. MSphere 3:e00148–00118.

Gloor, G. B., J. M. Macklaim, V. Pawlowsky-Glahn, and J. J. Egozcue. 2017. Microbiome datasets are compositional: and this is not optional. Frontiers in microbiology 8:2224.

Haines, D. F., J. A. Aylward, S. D. Frey, and K. A. Stinson. 2018. Regional patterns of floristic diversity and composition in forests invaded by garlic mustard (*Alliaria petiolata*). Northeastern Nat 25:399–417.

Hawkes, C. V., K. M. DeAngelis, and M. K. Firestone. 2007. Root interactions with soil microbial communities and processes. Pages 1–29 The Rhizosphere. Elsevier.

Hawkes, C. V., I. F. Wren, D. J. Herman, and M. K. Firestone. 2005. Plant invasion alters nitrogen cycling by modifying the soil nitrifying community. Ecology Letters 8:976–985.

Herlemann, D. P., M. Labrenz, K. Jürgens, S. Bertilsson, J. J. Waniek, and A. F. Andersson. 2011. Transitions in bacterial communities along the 2000 km salinity gradient of the Baltic Sea. ISME J 5:1571–1579.

Ihrmark, K., I. Bödeker, K. Cruz-Martinez, H. Friberg, A. Kubartova, J. Schenck, Y. Strid, J. Stenlid, M. Brandström-Durling, and K. E. Clemmensen. 2012. New primers to amplify the fungal ITS2 region–evaluation by 454-sequencing of artificial and natural communities. FEMS Microbiol Ecol 82:666–677.

Klindworth, A., E. Pruesse, T. Schweer, J. Peplies, C. Quast, M. Horn, and F. O. Glöckner. 2013. Evaluation of general 16S ribosomal RNA gene PCR primers for classical and next-generation sequencing-based diversity studies. Nucleic acids research 41:e1.

Klironomos, J. N. 2002. Feedback with soil biota contributes to plant rarity and invasiveness in communities. Nature 417:67–70.

Kõljalg, U., R. H. Nilsson, K. Abarenkov, L. Tedersoo, A. F. Taylor, M. Bahram, S. T. Bates, T. D. Bruns, J. Bengtsson-Palme, and T. M. Callaghan. 2013. Towards a unified paradigm for sequence-based identification of fungi. Wiley Online Library.

Lankau, R. A. 2011. Resistance and recovery of soil microbial communities in the face of *Alliaria petiolata* invasions. New Phytologist 189:536–548.

Lankau, R. A., V. Nuzzo, G. Spyreas, and A. S. Davis. 2009. Evolutionary limits ameliorate the negative impact of an invasive plant. PNAS 106:15362–15367.

Lee, M. R., S. L. Flory, R. P. Phillips, and J. P. Wright. 2018. Site conditions are more important than abundance for explaining plant invasion impacts on soil nitrogen cycling. Ecosphere 9:e02454.

Legendre, P., and E. D. Gallagher. 2001. Ecologically meaningful transformations for ordination of species data. Oecologia 129:271–280.

Li, W.-h., C.-b. Zhang, H.-b. Jiang, G.-r. Xin, and Z.-y. Yang. 2006. Changes in soil microbial community associated with invasion of the exotic weed, *Mikania micrantha* H.B.K. Plant Soil 281:309–324.

Nguyen, N. H., Z. Song, S. T. Bates, S. Branco, L. Tedersoo, J. Menke, J. S. Schilling, and P. G. Kennedy. 2016. FUNGuild: an open annotation tool for parsing fungal community datasets by ecological guild. Fungal Ecology 20:241–248.

Nuzzo, V. 1999. Invasion pattern of herb garlic mustard (*Alliaria petiolata*) in high quality forests. Biol Invasions 1:169–179.

Oksanen, J., F. G. Blanchet, R. Kindt, P. Legendre, R. O’hara, G. L. Simpson, P. Solymos, M. H. H. Stevens, and H. Wagner. 2010. Vegan: community ecology package. R package version 1.17-4. URL http://CRAN.R-project.org/package=vegan.

Parada, A. E., D. M. Needham, and J. A. Fuhrman. 2016. Every base matters: assessing small subunit rRNA primers for marine microbiomes with mock communities, time series and global field samples. Environmental microbiology 18:1403–1414.

Portier, E., W. L. Silver, and W. H. Yang. 2019. Invasive perennial forb effects on gross soil nitrogen cycling and nitrous oxide fluxes depend on phenology. Ecology 100:e02716.

Pringle, A., J. D. Bever, M. Gardes, J. L. Parrent, M. C. Rillig, and J. N. Klironomos. 2009. Mycorrhizal symbioses and plant invasions. Annu Rev Ecol Evol Syst 40:699–715.

Ramirez, K. S., J. W. Leff, A. Barberán, S. T. Bates, J. Betley, T. W. Crowther, E. F. Kelly, E. E. Oldfield, E. A. Shaw, and C. Steenbock. 2014. Biogeographic patterns in below-ground diversity in New York City’s Central Park are similar to those observed globally. Proceedings of the royal society B: biological sciences 281:20141988.

Roberts, K. J., and R. C. Anderson. 2001. Effect of garlic mustard [*Alliaria petiolata* (Beib. Cavara & Grande)] extracts on plants and arbuscular mycorrhizal (AM) fungi. Am Midl Nat 146:146–152.

Rodgers, V. L., K. A. Stinson, and A. C. Finzi. 2008. Ready or not, garlic mustard is moving in: *Alliaria petiolata* as a member of eastern North American forests. Bioscience 58:426–436.

Saikkonen, K. 2007. Forest structure and fungal endophytes. Fung Biol Rev 21:67–74.

Schoch, C. L., K. A. Seifert, S. Huhndorf, V. Robert, J. L. Spouge, C. A. Levesque, W. Chen, and F. B. Consortium. 2012. Nuclear ribosomal internal transcribed spacer (ITS) region as a universal DNA barcode marker for Fungi. PNAS 109:6241–6246.

Smith, G. R., L. C. Edy, and K. G. Peay. 2021. Contrasting fungal responses to wildfire across different ecosystem types. Molecular Ecology 30:844–854.

Smith, L. M., and H. L. Reynolds. 2015. Extended leaf phenology, allelopathy, and inter-population variation influence invasion success of an understory forest herb. Biological invasions 17:2299–2313.

Stinson, K. A., S. A. Campbell, J. R. Powell, B. E. Wolfe, R. M. Callaway, G. C. Thelen, S. G. Hallett, D. Prati, and J. N. Klironomos. 2006. Invasive plant suppresses the growth of native tree seedlings by disrupting belowground mutualisms. PLoS Biol 4:727–731.

Team, R. C. 2013. R: A language and environment for statistical computing.

Urbanowicz, C., V. J. Pasquarella, and K. A. Stinson. 2019. Differences in landscape drivers of garlic mustard invasion within and across ecoregions. Biological invasions 21:1249–1258.

Vaughn, S. F., and M. A. Berhow. 1999. Allelochemicals isolated from tissues of the invasive weed garlic mustard (*Alliaria petiolata*). J Chem Ecol 25:2495–2504.

Wang, Q., G. M. Garrity, J. M. Tiedje, and J. R. Cole. 2007. Naive Bayesian classifier for rapid assignment of rRNA sequences into the new bacterial taxonomy. Applied and Environmental Microbiology 73:5261–5267.

White, T. J., T. Bruns, S. Lee, and J. Taylor. 1990. Amplification and direct sequencing of fungal ribosomal RNA genes for phylogenetics. PCR protocols: a guide to methods and applications 18:315–322.

Wolfe, B. E., V. L. Rodgers, K. A. Stinson, and A. Pringle. 2008. The invasive plant *Alliaria petiolata* (garlic mustard) inhibits ectomycorrhizal fungi in its introduced range. J Ecology 96:777–783.

